# Human reproduction comes at the expense of faster aging and a shorter life

**DOI:** 10.1101/2024.07.18.603826

**Authors:** Mikaela Hukkanen, Anna Kankaanpää, Aino Heikkinen, Jaakko Kaprio, Robin Cristofari, Miina Ollikainen

**Affiliations:** Institute for Molecular Medicine Finland, University of Helsinki, Tukholmankatu 8, 00290 Helsinki, Finland; Minerva Foundation Institute for Medical Research, Tukholmankatu 8, 00290 Helsinki, Finland; Gerontology Research Center (GEREC), Faculty of Sport and Health Sciences, University of Jyväskylä, Rautpohjankatu 8, 40014, Jyväskylä, Finland; Institute of Biotechnology, Helsinki Institute of Life Science, University of Helsinki, Viikinkaari 5, 00790, Helsinki, Finland

**Keywords:** Reproduction, resource allocation, epigenetics, survival, biological aging, twin cohorts

## Abstract

Evolutionary theories suggest a trade-off between resources allocated to reproduction and those allocated to self-maintenance, and predict that higher reproductive output entails a shorter lifespan. This study investigates the impact of childbearing on aging and lifespan using data from contemporary Finnish twin women. We model the association between reproductive trajectories and survival in 17,080 women, and assess biological aging in a subset of participants (N=1,117) using the PCGrimAge clock, an algorithm trained to predict biological aging and mortality risk from DNA methylation. Our findings suggest that early childbearing, numerous pregnancies or nulliparity all contribute to accelerated aging and increased mortality risk. These results provide strong evidence for the existence of a trade-off between reproduction, aging and lifespan in modern humans, and provide novel insights into the genetic and lifestyle determinants of healthspan.

---

**Reproductive success and lifespan determine fitness, and the resource allocation trade-off between these two functions is a key tenet of evolutionary biology**^**1,2**^. **Previous studies support such a trade-off by suggesting that an increased number of offspring can reduce healthspan across and within species**^**3–5**^, **but it remains unclear whether (i) increased reproductive investment accelerates the aging process itself, and (ii) what role do genetics, lifestyle, and the timing of reproduction play in this acceleration. This study investigates whether pregnancy, one important cost of reproductive investment**^**6**^, **influences aging and lifespan in contemporary humans independent of lifestyle and genetic determinants. By examining the number and timing of pregnancies in a large cohort of Finnish twins**^**7**^, **we identified reproductive trajectories that predict both lifespan and DNA methylation-based biological aging**^**8,9**^. **We show that early pregnancy and a higher number of pregnancies throughout life are associated with accelerated epigenetic aging and reduced lifespan: interestingly, at the other end of the spectrum, nulliparous women also experience similar effects. We used a powerful twin design to determine the familial contribution to lifespan and aging, and revealed a strong genetic influence on these traits. Our findings bring unambiguous support to evolutionary theories of aging**^**1**^, **and contribute to the rapidly growing field of evolutionary medicine**^**10**^, **providing novel insights into the determinants of healthy lifespan**.

Within the lifespan of an individual organism, the resource-allocation theory of aging supposes that resources invested into reproduction are taken away from resources allocated to survival, through e.g., cellular maintenance mechanisms^1-3^. Consequently, aging — understood here as the gradual decline in physical capabilities over time — is proposed to have evolved from selection favoring traits that promote reproductive success^1^. Multiple studies have found a U-shaped relationship between reproduction and mortality in humans and other species: both nulliparity and giving birth to many offspring has been found to increase mortality risk^4,5,11^. At one end of the spectrum, lower survival prospects and low breeding output may both be caused by poorer initial health - but at the other end of the spectrum, decreased lifespan and high breeding output may reveal the costs of increased reproductive investment, as predicted by life-history theory^5^. However, lifespan is only a proxy for the aging process itself, and can be influenced by external events: as a consequence, it is still unknown whether increased reproductive output really accelerates the aging process, and, if yes, whether this acceleration is already observable before old age.

DNA methylation-derived age estimates — the epigenetic clocks — can be used to approximate one’s biological age as the apparent age of the body^12^. Further, the difference between known chronological age, and estimated biological age, gives us access to an individual’s aging rate, i.e. the relative speed of each individual’s own clock^13^. Epigenetic clocks can be categorized based on their training focus: those trained to estimate chronological age as closely as possible, such as Horvath’s clock^14^ and Hannum’s clock^15^, and those trained to assess frailty or “phenotypic” age, including the DunedinPACE^16^, PhenoAge^17^ and GrimAge^8^ clocks. Specifically, DunedinPACE is trained to measure the pace of aging, using longitudinal data to capture the rate of biological deterioration over time, while PhenoAge and GrimAge are trained to reflect biological age by incorporating clinical biomarkers and mortality risk. The GrimAge epigenetic clock, in particular, excels in predicting time-to-death, offering a valuable measure of frailty, years before the end of life^8^. A recently developed principal component (PC)-based version of the GrimAge clock (hereafter “PCGrimAge”), uses PCs derived from CpG-level DNA methylation data for biological age estimation^9^. The PCGrimAge has enhanced the reliability of biological age estimates, as shown by reduced deviations between replicates compared with the original CpG-level clock^9^. The present study is based primarily on that clock.

Some pioneering cross-sectional studies^3,18,19^ have linked the number of pregnancies to epigenetic age acceleration, but the evidence has been equivocal^20^. Recently, Ryan et al. (2024)^3^ found that increasing parity accelerated epigenetic aging in 22 year old women using six independent clocks. However, like in other previous studies, the focus on younger women without assessing their survival prospects severely limits our understanding of the consequences of reproduction on ageing. What is more, all studies to our knowledge have focused on the number of children or the age at first birth, regardless of the timing of subsequent pregnancies, and thus could not consider the effects of the full childbearing history, including later-life pregnancies. Finally, it has so far been challenging to assess the influence of confounding factors, and first and foremost of shared genetic determinants (e.g. as a genetic control of pace-of-life^21^), on the correlation between pregnancy and aging. Thus, the long-term consequences on epigenetic aging trajectories and survival of lifelong reproductive history on the one hand, and familial and lifestyle factors on the other remain largely unknown.

The current study addresses this knowledge gap by investigating whether pregnancy influences aging and lifespan in modern day humans. Based on ⪆17,000 participants of the Finnish Twin Cohorts^7^, our approach accounts for both the number and the timing of child births by identifying underlying typical reproductive trajectories through latent class analysis^22^. If indeed reproduction comes at the expense of somatic maintenance, we expect the women with the trajectories of higher lifetime reproductive output to display accelerated epigenetic aging and shorter lifespan. We test this by modeling survival as a function of the reproductive trajectories. Among those women with methylation data (N⪆1000), we assess differences in biological age acceleration using the PCGrimAge^8,9^ algorithm. By studying monozygotic and dizygotic twins, we are able to adjust for the effect of confounding familial background including genetics. Our method also incorporates data on socioeconomic background and lifestyle, adjusting for prevalent risk-factors of later life morbidity of the 21st century, allowing a rigorous examination of the interplay between reproduction, aging and lifespan. Our results suggest that early childbearing and increased number of lifetime pregnancies accelerates aging and increases mortality risk compared with women who have only a few pregnancies later in life.

## Results and discussion

### Reproductive classes

In order to account not only for the number but also for the timing of pregnancies, we first modelled individual reproductive trajectories as a mixture of K typical trajectories inferred from lifelong pregnancy data (see Methods and Supplementary Methods, Supplementary Results) using latent class analysis (LCA^22^) in Mplus version 8.2^23^. This data included 10,783 women, who had had at least one pregnancy during their life. We tested models with live births summed over 1 to 4 years bins, and K=1 to 8 latent classes, and selected the one minimizing sample-size adjusted BIC, with the constraint of keeping a meaningful sample size (at least 3% of women to be most likely in the smallest class), and maximizing entropy and class-specific posterior probabilities. In the selected model, individual reproductive trajectories were described as a mixture of K=6 classes (hereafter “reproductive classes”), where live births were summed over 3-year age bins (see Methods and Suppl. Methods; latent class parameters are given in Table 1, cross-class classification probabilities are given in Extended Data Table 1). In this framework, each reproductive class is thus defined as a typical timing and number of children (Fig. 1A): for example, a woman in class 1 had, on average, two children (SD=1) and a high probability to give birth before age 24, while a woman in class 6 would have many children (6.8, SD=2.4) interspersed throughout her reproductive life. This result was well supported by parametric bootstrapping (Extended Data Fig. 1); under this model, individuals are mostly assigned to a single class (see number of children by class in Extended Data Figure 2). Women who did not have children (N=6,392) were added to this data and assigned to a separate class (class 0 i.e. “nulliparous”) with a posterior probability of ∼1 for further analyses. The ability of our model to robustly and convergently identify these classes suggests that contemporary Finnish women can indeed be distinguished by their reproductive histories into distinct subgroups. This might indicate that modern humans exhibit, for instance, varying underlying behavioral and cultural traits^24^, as well as genetic liability^25^ that influence their family-starting decisions and reproductive behaviors.

**Table 1.**
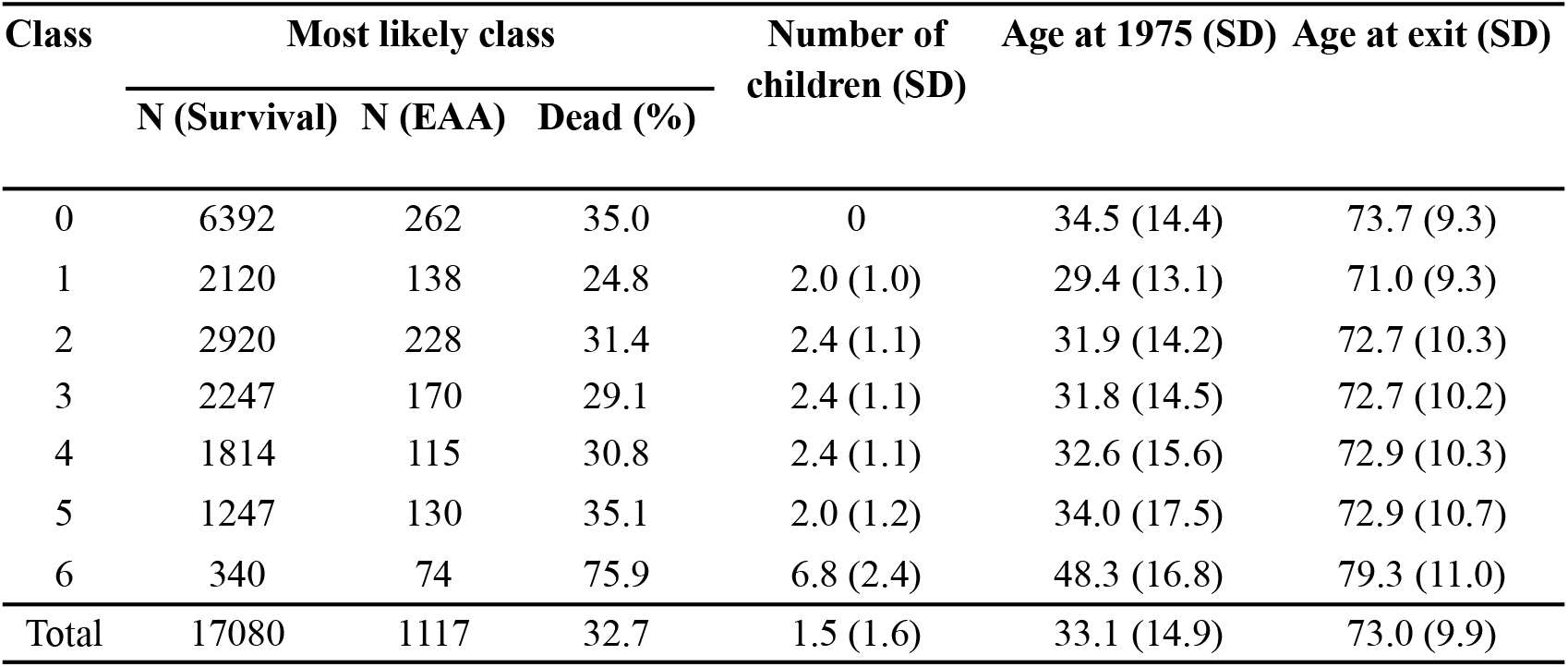
Latent class characteristics. Number of women in survival analysis and epigenetic aging analysis (EAA), and the proportion of dead at the end of the follow-up for each class when women were assigned to the most likely class based on highest posterior probabilities from the latent class solution. The number of children, age at the start of the follow-up in 1975 and age at exit (death or end of the follow-up) with their respective standard deviations (SD) are given for each class, calculated as weighted averages and standard errors based on class-specific posterior probabilities. The bottom row indicates sample sizes, averages and standard errors across all classes.

**Figure 1.**
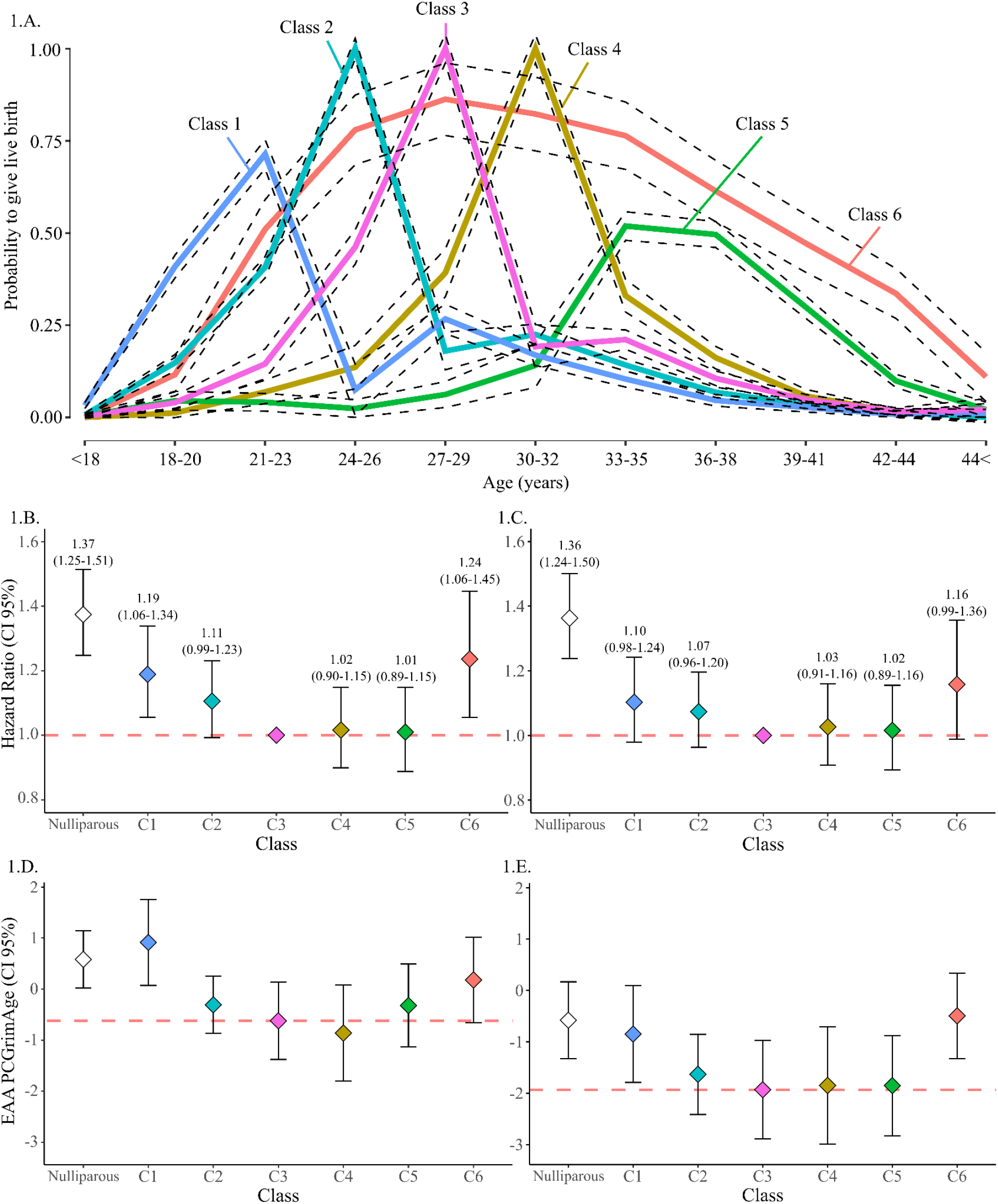
Reproductive classes and associations with survival and epigenetic age acceleration (EAA). A. Latent class model with 6 classes and 3-year age bins with 95% confidence intervals. B. Mortality hazard ratios with 95% confidence intervals for each latent class from a Cox proportional hazards model. The dashed red line indicates the reference class used in the survival analysis (class 3). C. Mortality hazard ratios with 95% confidence intervals for the fully adjusted model with additional covariates for lifestyle factors. The dashed red line indicates the reference class used in the survival analysis (class 3). D. Estimates and 95% confidence intervals for EAA by the PCGrimAge clock for each class. The dashed red line is drawn for comparison to survival analysis at the estimates for class 3. E. Estimates and 95% confidence intervals for EAA by the PCGrimAge clock for each class from a weighted Bolck-Croon-Hagenaars model, with additional adjustments for lifestyle factors. The dashed red line is drawn for comparison to survival analysis at the estimates for class 3.

### Reproductive classes predict survival

Survival differed significantly between reproductive classes when modeled as mixed Cox proportional hazards, weighted on probabilities of belonging to each class (N=17,080, see Methods, Supplementary Methods and Fig. 1B-C). When accounting for historical birth cohort and relatedness, women with no pregnancies, or with few pregnancies early in life (classes 0, 1 and to a lesser extent 2) had increased risk of death compared to the reference class i.e. class 3 (hazard ratios (HR)=1.37, 1.19, and 1.11, respectively, see Fig. 1A-1B). Symmetrically, women with many children throughout life (class 6) also had increased mortality (HR=1.24, see Fig. 1B). This pattern was attenuated but not abolished when correcting for known confounding factors (BMI, tobacco and alcohol use, and education, see Methods, Supplementary Methods and Fig. 1C), also for the complete case sample with no covariate imputation (Extended Data Figure 3A, see Supplementary Methods for details).

### Reproductive classes predict aging

Similarly to survival, there were significant differences between the reproductive classes in epigenetic aging measured by the PCGrimAge clock in a subsample of 1,117 women (Figure 1D-E, see comparisons between all classes in Extended Data Table 4). In our minimally adjusted model (controlling for chronological age and relatedness, see Methods and Supplementary Methods), we observed the most accelerated aging for women with few pregnancies in early life (class 1), with 1.77 years (SE=0.65) higher biological aging rate than class 4, that had the lowest aging rate. Nulliparous women (class 0) and women with many pregnancies throughout life (class 6) also showed more accelerated aging compared with the women having only a few children in their late 20s (classes 2-4). When adjusting for lifestyle factors (BMI, tobacco and alcohol use, and education), the most accelerated aging was observed for the class with the highest reproductive investment (class 6) with 1.44 (0.63) years higher epigenetic aging rate than the lowest class 3. These results suggest that the effect of reproductive investment on aging is not a mere consequence of confounding lifestyle factors, for example, the people with many children also living a healthier life for e.g. cultural or religious reasons. The observed patterns were also robust to the method of handling missing covariate data, as the adjusted model resulted in similar observations using multiple imputation versus full-information maximum likelihood methods (Figure 1D vs Extended Data Figure 3B). We tested whether we would observe similar patterns in two other published epigenetic clocks that are trained to capture different aspects of biological aging: DunedinPACE^16^, which is trained to predict pace of biological aging using 19 indicators of organ-system integrity, and PC-based^9^ PhenoAge^17^, a clock that is developed to measure “phenotypic age” from clinical measures^AH^. We observed similar patterns with the DunedinPACE estimator as for PCGrimAge, yet not for PCPhenoAge, likely reflecting the different calibration strategies and training populations used when building these clocks (Extended Data Figure 4).

### Links between reproduction and longevity

The observed pattern of increased mortality risk and accelerated aging in nulliparous women (class 0), mothers of early childbirth (class 1), and high lifetime parity (class 6) are in line with previous findings on the U-shaped pattern between parity and mortality risk, and the effects of early motherhood on later health^11,26,27^. Among the nulliparous women, pre-existing poor health, or high lifestyle risks, might make reproduction less likely, confounding the observed increased mortality risk and accelerated aging^26,28^. However, the high mortality risk of nulliparous women may also be influenced by the protective effects of pregnancy and lactation towards certain diseases, like hormonal cancers, and by the lack of social support from children^26,28,29^. Interestingly, women who reproduce early also have increased mortality risk and accelerated aging. This may be partly explained by an unfavourable socioeconomic background and having limited access to healthcare and resources in general^26,30^. Further, early childbirth has been associated with increased risk for later life obesity and mobility impairment, and with low education, restricted occupational progression and high risk of divorce^31,32^. Simultaneously, higher education is known to be associated with healthier lifestyle and increased survival while delaying pregnancy, likely explaining the longevity and slowest aging in the classes of later motherhood^33^. While these factors related to socioeconomic status and health are partially accounted for through the inclusion of known lifestyle descriptors in the current study, they are still likely to contribute to our findings. Finally, childbearing itself is expected to be a major determinant of survival in the high lifetime reproduction group, as predicted by life-history theory^1^. Pregnancy, childbirth, and lactation pose significant physiological challenges^6^, especially for young mothers^34^. Early mothers may also be less resilient to parenting-related physical, emotional, and economic stress^35^, while experiencing higher cumulative stress and allostatic load overall^30^. At the epigenomic level, the accelerated aging in the class of highest reproductive investment may result from pregnancy-induced changes in for example, i) CpG-specific methylation of genes involved in growth and differentiation^18,36^, ii) epigenetic stability due to oxidative stress^37^, iii) or blood cell type composition due to immune activation^38^.

### Familial effects on survival and aging

The variance explained by the familial background for survival between the two twins from the same pair were 0.66 and 0.26 for monozygotic (MZ) and dizygotic (DZ), respectively. This suggests a strong genetic influence in survival, as MZ twins in a pair are identical at the genome sequence level, while DZ twins share on average only half of their segregating genes by descent from their common parents (Methods, Supplementary Methods). The pronounced genetic effect echoes previous estimates of the heritability of human lifespan^39^. For epigenetic age acceleration (EAA) PCGrimAge, the intra class correlation coefficient (ICCs) for MZ and DZ twins were 0.619 and 0.387, respectively (Methods, Supplementary Methods). The heritability of epigenetic aging was estimated at 47% (CI 15.0-77.8%), indicating a more pronounced genetic component than for survival, replicating previous findings^14,40–42^.

## Conclusions

To our knowledge this is the first study to report distinct reproductive trajectories of pregnancy history in contemporary women. Supported by life-history theory, these classes were predictive of survival, as the women belonging to the classes of nulliparity, early motherhood and extreme lifetime reproductive investment had increased mortality risk. For the first time, we were able to link these differences in reproductive history and survival to biological aging, years before death. Furthermore, by utilizing a powerful twin design we were able to determine the contribution of familial background on lifespan and aging, revealing a strong genetic influence on these traits, replicating previous findings and providing novel insight into the determinants of healthy lifespan.

## Methods

### Study participants and anthropometric data

The 17,175 study participant women were a subset of the older Finnish Twin Cohort^7^ born between 1880 and 1958, with both co-twins alive in 1967 (see Supplementary Methods, Supplementary Results and Extended Data Table 3). Data on the number and birth year of children was collected in a mailed questionnaire in 1975 and validated from the Digital and Population Data Services Agency for the women born after 1949 (see Supplementary Methods). This data was used as a proxy for pregnancy history. Information on major predictors of mortality and hence potential confounders (education, height, weight, smoking and alcohol use) were collected in the mailed questionnaire, or imputed in case of missingness using Multiple Imputation^43^ in Mplus^23^ (version 8.2), assuming that these values are missing at random^43^ (see Supplementary Methods and Extended Data Table 3). These data were used to compile covariates in survival and EAA analyses, described in detail in Supplementary Methods. Briefly, year of birth was used to subset the participants into age groups to compile a categorical variable describing the cohort experience of living circumstances. Education was included in the models as a three-level categorical variable (primary ≤7 years of lifetime education, secondary ≤10 years, tertiary >10 years), body-mass index (BMI) was modeled as a continuous variable (kg/m^2^) with a centered and a quadratic term, whereas categorical variables were used to model smoking status (never, former, infrequent, light, medium, heavy) and alcohol use (lifetime abstainer, former, infrequent, low, medium ≤). Time to death was retrieved from the Population Information Services of the Digital and Population Data Services Agency.

### Reproductive trajectory analysis

Reproductive trajectories were identified from the live birth data of parous women (10,783) using latent class analysis (LCA)^22^ with the Mplus software, using maximum likelihood estimators with robust standard errors (MLR) using a sandwich estimator^44^ (Supplementary Methods). We tested models summing the number of live births over 1-4 year bins, and testing models with 1 to 8 latent classes. The optimal model was determined by minimizing Akaike’s information criterion (AIC)^45^ and sample size adjusted Bayesian information criterion (saBIC)^46,47^, while aiming to maximize the relative sample size of the smallest class, entropy (a measure of class separation) and class-specific posterior probabilities. See Supplementary Methods and Extended Data Table 2 for model comparison including full details. The final model consisted of 6 classes of reproductive trajectories by 3-year age bins, where the nulliparous were added as a 7th class. The robustness of the 6 class 3-year solution was tested with parametric bootstrapping (see Supplementary Methods).

### Survival analysis

We used a mixed Cox proportional hazards model in R with packages *survival*^48^ and *coxme*^49^ to model survival in all women that were confirmed to be twins and lived past 40 years (N=17,080). We used proportional assignment^50^ to reflect uncertainty in the classification using class-specific posterior probabilities as weights. All models were controlled for non-independence among twins from the same pair using twin pair ID as a random effect, left truncation and right censoring. Two models were fitted: 1) unadjusted model with historical birth cohort and twin pair ID variables, and 2) adjusted model with additionally smoking, alcohol use, and BMI in 1975, and lifetime education as independent variables. For the adjusted model, we report the results using the imputed covariates in the Main Text, and the results using the data with no imputation in Extended Data Figure 3. See full details of modeling the differences in survival between the latent classes including the nulliparous class in Supplementary Methods.

### DNA methylation assay and Epigenetic Age Acceleration

Epigenetic age was assessed for a subsample of the participants (N=1,117) based on peripheral blood samples taken at an age ranging from 38 to 86^7^. DNA methylation levels were measured with Illumina’s Infinium HumanMethylation450 BeadChips (450k, N=250) or the Infinium MethylationEPIC BeadChips versions 1 and 2 (EPICv1, N=667, and EPICv2, N=200). Quality control and control probe-based quantile normalization were conducted using R^51^ (version 4.2.3) with package *meffil*^52^. Beta Mixture Quantile Normalization^53^ to adjust the beta-values of type II design probes into a statistical distribution characteristic of type I probes was conducted using the package *wateRmelon*^54^. See Supplementary Methods for quantifying and preprocessing DNA methylation data for epigenetic aging.

Biological age was determined from methylation beta values using principal component (PC)-based^9^ GrimAge^8^ (“PCGrimAge”, reported in the Main Text) and PhenoAge^17^ (“PCPhenoAge”) as well as DunedinPACE^16^ (reported in the Extended Data). Epigenetic age acceleration was determined as the residual of epigenetic age linearly regressed against chronological age in our sample of 1,117 participants.

The differences in epigenetic age acceleration were modeled in MPlus using the Bolck-Croon-Hagenaars (BCH) approach^55^ (see Supplementary Methods for full details). We used class-specific weights as training data to model epigenetic age acceleration between the latent classes and the nulliparous women by fitting two models: 1) the unadjusted model included independent variables for the twin pair ID, chronological age and historical cohort, while 2) the adjusted model included additional variables for smoking, alcohol use, BMI at the time of blood sampling, and lifetime education. As for the survival models, we report the adjusted model results using the imputed covariates in the Main Text, and the results using the data with no imputation in Extended Data Figure 3.

### Familial effects

The variance explained by the familial background was reported as the variance explained by twin pair IDs in the mixed Cox models with no explanatory variables, adjusted for left truncation. The variation explained by familial background in the epigenetic age acceleration was reported as the ICC of the PCGrimAge estimates. Heritability was defined as: 2x(ICC_MZ_-ICC_DZ_). The confidence intervals for epigenetic aging were calculated from the standard errors of the ICCs: 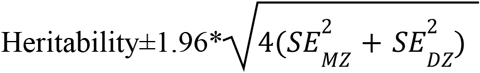, where SE indicates the standard errors of the ICCs for MZ and DZ twins separately.

## End notes

## Acknowledgements

The authors want to acknowledge the computational resources of the Institute of Molecular Medicine Finland (FIMM) Technology Center. Funding. This research was supported by funding from the Emil Aaltonen Foundation to Mikaela Hukkanen (#230051), the University of Helsinki Doctoral School of Population Health (DOCPOP) to Aino Heikkinen, the Academy of Finland Center of Excellence in Complex Disease Genetics to Jaakko Kaprio (#352792), the Research Council of Finland (#328685, #307339, #297908 and #251316 to Miina Ollikainen, and #331320 and #354649 to Robin Cristofari), and the Minerva Foundation, the Liv o Hälsa sr., and the Sigrid Juselius Foundation to Miina Ollikainen.

## Author contributions

The study was initiated and designed by Mikaela Hukkanen, Jaakko Kaprio, Robin Cristofari and Miina Ollikainen. Data acquisition and preparation were performed by Jaakko Kaprio, Miina Ollikainen, Mikaela Hukkanen and Aino Heikkinen. Analysis of the data was conducted by Mikaela Hukkanen and Anna Kankaanpää. The first draft of the manuscript was written by Mikaela Hukkanen, and all authors commented on the manuscript. All authors read and approved the final manuscript.

## Additional Information

### Supplementary Information statement

Supplementary Information (Methods and Results) are available for this paper.

### Ethics approval

This study has been approved by the appropriate national research ethics committees, with the most recent approval granted by the Hospital District of Helsinki and Uusimaa ethics board in 2018 (#1799/2017). Blood samples for DNA analyses were collected from each participant after they had signed a written informed consent.

