## Supplementary Information for "Human reproduction comes at the expense of faster aging and a shorter life"

### Supplementary Methods

#### 1. Participants

The study participants were a subset of the older Finnish Twin Cohort (FTC)[7], a population-based prospective cohort consisting of twins from monozygotic (MZ) and dizygotic (DZ) pairs born before 1958 with both co-twins. A mailed questionnaire survey was sent to all twin pairs whenever both of the twins were alive in 1967[7]. The questionnaire data includes self-reported information on for example family members, education, weight, height, and lifestyle-related factors such as smoking and alcohol consumption. The current study includes 17,175 women from the cohort, who were born between 1880 and 1957, and had a known number of children. The full sample was used to compile the reproductive trajectories in latent class analysis, while a sample of 17,080 women who lived at least 40 years and were verified as twins was used to assess the link between reproductive history and mortality risk. For a subsample of 1,117 participants, blood samples were taken during 1994-2020. These samples were used to assess genome-wide DNA methylation and epigenetic age acceleration.

#### 2. Data

##### 2.1. Live birth data

Live birth data was curated for all the study participants from the 1975 questionnaire. Participants were requested to report the birth years of each son and daughter. In case a woman had reported multiple children within the same calendar year, the children were considered to be from a multiple pregnancy (i.e. twins, triplets, etc.). This data was validated for all participants born between 1950-1957 (N=4,431, 41.1%) by retrieving the exact birth dates of each child from the Digital and Population Data Services Agency on 31 December 2009.

#### 2.2. Contextual data

We adjusted our statistical analyses with variables that were hypothesized to contribute to variance in epigenetic age acceleration and mortality risk. See sample sizes by variable categories in Extended Data Table 3.

*Historical birth cohort*

Our study participants were born between 1880 and 1957, a period during which Finland went through drastic societal and political changes, with likely influence on the study participants' health and survival prospects. To account for these differences in living circumstances, we assigned each participant to their birth cohort, reflecting differing living circumstances in each time period: 1880-1898 Industrial era, 1899-1917 Pre-independence, 1918-1938 Post-independence, 1939-1945 World War II, 1946-1957 Cold War.

*BMI*

Body mass index (BMI, $kg/m^2$) has been shown to be associated with accelerated epigenetic aging and increased mortality risk[56,57]. For survival analysis, BMI was calculated from self-reported weight and height in 1975, which have been previously shown to correlate well (r(Pearson)=0.95, CI=0.94-0.96) with measured values in the same cohort[58]. For epigenetic aging analysis, BMI at the time of blood-draw was used. As BMI may have a non-monotonous relationship with health[59], we used centered BMI as a linear term as well as a quadratic term in both survival and epigenetic aging models.

*Smoking*

Smoking is also a known risk factor for mortality and has been shown to influence DNA methylation and accelerate epigenetic aging[56,60]. For survival analysis, participants were attributed a smoking status ("never", "former", "infrequent", "light" (1-9 cigarettes per day), "medium" (10-19 cigarettes per day), "heavy" 20< cigarettes per day) based on questionnaire data from 1975[61]. At the time of blood sampling, the participants were asked a three-level smoking status ("never", "former", "current"), which was used as a covariate in the epigenetic aging analysis.

*Alcohol*

Alcohol use is associated with DNA methylation, epigenetic aging, as well as morbidity and mortality risk[62,63]. An alcohol use index was compiled from each questionnaire, first as a continuous variable reflecting average daily use in grams of ethanol[61], further binned into a categorical variable ("abstainer", "infrequent" (<1.3g/day), "low" (<25g/day), "medium" (<45g/day), "high" (<65g/day), "higher" (≥65g/day) as described in Zhao et al. (2023)[64]. As heavy drinking was rare among the women in our cohort, the "medium", "high", and "higher" categories were combined into a "medium or high" category in the survival analysis. In the epigenetic aging analysis, the alcohol index at the time of blood sampling was binned to three levels ("non-drinker"=0g/day, "infrequent"<1.3g/day, "frequent"≥1.3g/day) due to the relatively small sample sizes for different categories within each reproductive class.

*Education*

Higher education has been shown to be associated with lower EAA, later-life morbidity and delayed mortality[65,66]. Education is also hypothesized to reflect differences in socio-economic status and social networks[67]. Lifetime years of education were compiled from answers about attained educational level in 1975 and 1981, as described in Silventoinen et al. (2004)[68.]. We used education as a three-level ("primary", "secondary", "tertiary") categorical variable in our analysis for both mortality and epigenetic aging.

##### 2.3. Handling of missing data

As not all participants answered the 1975 questionnaire (Extended Data Table 3), we used Multiple Imputation (MI) to handle missing covariate values using Mplus[AX] software (version 8.2), assuming that these values are missing at random[43] (alcohol use, smoking, BMI and education). MI is assumed to produce unbiased parameter estimates and standard errors, as all the available data is utilized to estimate missing variables[43]. In both survival and epigenetic aging models, 20 datasets (as recommended in Little et al. 2013[43]) with imputed covariates were generated and the subsequent modeling was conducted in the same way as the complete case models, by handling multiple imputed datasets with the R package *mitools*[69]. For comparison, we also modeled the final analyses with the complete case samples without imputation for both, survival and epigenetic aging (Extended Data Figure 3).

##### 2.4. Epigenetic aging data

*Blood sampling, DNA extraction, beadchips*

Epigenetic age was assessed for a subsample of the participants (N=1,117) based on peripheral blood samples taken at an age ranging from 36 to 89[7]. High molecular weight DNA was extracted from the blood samples with standard automated protocols and bisulfite converted using EZ-96 DNA/methylation-Gold Kit (Zymo Research, Irvine, CA, USA) according to the manufacturer's protocol. DNA methylation levels were measured using Illumina's Infinium HumanMethylation450 BeadChips (450k) or the Infinium MethylationEPIC BeadChips versions 1 and 2 (EPICv1 and EPICv2), which quantify methylation levels at single-nucleotide resolution of over 450,000, 850,000 and 935,000 CpG-sites, respectively. Of the samples included in the present study, 250 were assayed using the 450k array, and 667 and 200 using the EPICv1 and EPICv2 arrays. Both twins in a pair were assayed with the same platform.

*Preprocessing*

DNA methylation data was preprocessed in R[51] keeping data from each array type separate. Quality control and control probe-based quantile normalization were conducted using the R package *meffil*[52]. We validated our dataset by ascertaining that all methylation-based predictors of sex were indeed predicting female sex. We discarded samples where 1) median methylated signal over all CpG sites was more than 3 SD from the expected, based on regression of median methylated signal by median unmethylated signal in all samples, 2) BeadChip inherent control probe values deviated over 5 SD from from the overall control probe mean, 3) over 20% of probes had detection p-value of over 0.01, and 4) over 20% of their probes had less than 3 detected beads. Additionally, a CpG site was removed in all samples if 1) there was only background signal in over 20% of the samples (detection p-value > 0.05), and 2) if the bead count of a certain probe was less than 3 in over 20% of samples. Further, all probes in sex chromosomes as well as cross-reactive probes and polymorphic CpGs were removed[70,71]. Raw probe intensities for each sample were adjusted to conform to its set of normalized quantiles based on 16, 15 and 16 first control probe principal components, for 450k, EPICv1 and EPICv2 platforms respectively (Extended Data Figure 5). Beta Mixture Quantile Normalization[53] to adjust the beta-values of type II design probes into a statistical distribution characteristic of type I probes was conducted using the package *wateRmelon*[54]. Methylation beta values were obtained by dividing the intensity of methylated sites (M) by the sum of intensities from methylated and unmethylated sites (Beta value = M / (M + U), where U represents the intensity of unmethylated probes). These processed beta values were then used as input for calculating epigenetic aging.

*Epigenetic aging*

Biological age was determined from DNA methylation beta values using three published algorithms: principal component versions[9] of GrimAge[8] and PhenoAge[17], as well as DunedinPACE[16]. Epigenetic ages of the above algorithms were calculated in R using the scripts provided in the corresponding publications. Epigenetic age acceleration was determined as the residual of epigenetic age linearly regressed against chronological age in our sample of 1,117 participants.

##### 2.5. Mortality

The mortality follow-up started in 1975 and ended on 31 December 2020, when dates of death were retrieved from the Digital and Population Data Services Agency Finland. The survival time for each woman was defined in years until the date of emigration, death, or end of follow-up, whichever came first.

#### 3. Statistical analysis

##### 3.1. Reproductive trajectories

*Latent class analysis*

Reproductive trajectories were identified from lifelong live birth data of 10,783 women using latent class analysis (LCA)[22] with the Mplus software. The process iterates from random starting values to explore different regions of the parameter space and find values that maximize the likelihood of observing the data given the model. The estimated parameters include class-specific item response probabilities (to give birth) for each indicator (ages at childbirth, see below), as well as class probability parameters, which are used to calculate individual-level posterior probabilities of belonging to each class. The estimation was done by using maximum likelihood estimators with robust standard errors (MLR) using a sandwich estimator (MLR type=COMPLEX)[44]. The sandwich estimator was used as it is robust to violations of non-independence among observations resulting from shared inheritance. For initiating the estimation we used 500 random sets of parameter values and 20 final stage optimization rounds; the number of starting values and optimizations was increased when needed for the model to converge to global maximum[72]. Global maximum was assumed when the final models converged to the same likelihood at the final stage optimization rounds[72].

The optimal model solution was determined following guidelines by Sinha et al. (2021)[73], suggesting to select the model with fewest number of classes that best fits the data. Fit was assessed with Akaike's information criterion (AIC)[45] and sample size adjusted Bayesian information criterion (saBIC)[46,47], which both assess model accuracy based on likelihood but penalize overfitting[73]. If these criteria disagreed we prioritized saBIC as it is recommended especially for analyses where sample size is large[74,75]. After prioritizing information criterion, we aimed to maximize the relative sample size of the smallest class, entropy (a measure of class separation) and class-specific posterior probabilities. We tested models with number of live births summed over 1-4 year bins as the indicator variables (for example in the 3-year age bins, all births younger than 18 years, 18-20 years, 21-23 years, … , older than 44 years). For each age bin, we fitted latent class models with 1 to 8 classes. The number of live births within 3-year age bins were selected as indicator variables for the final latent class analysis, with 6 latent classes. Comparisons between all the tested models can be found in Extended Data Table 2. In the final model solution, we added people with no reported live births i.e. the nulliparous as an additional class, for which an individual was assigned a probability of 0.9999999 if they had no live births (and 0.0000001 as the class-specific posterior probabilities), and 0.0000001, if they had given birth during the study period and were included in the LCA solution.

*Parametric bootstrapping*

The robustness of the final LCA model was tested with parametric bootstrapping. We simulated 100 datasets consisting of 10,783 individuals. For each age bin $i$, the individual probability of having $n$ children was calculated as the product of the posterior class belonging probability matrix (I classes x S individuals), by the latent class matrix (N possible children x I classes), yielding a class-specific N possible children x S individuals) probability matrix. For each bootstrap replicate, an actual number of children in each age class for each individual was drawn randomly according to these probability matrices. For each simulated dataset, the latent class model with 6 classes was then fitted.

##### 3.2. Survival

The differences in survival between the latent classes including the nulliparous class were modeled in R. To account for the non-independence between women from the same twin pair, we conducted a mixed Cox proportional hazards model, where twin pair ID was used as a random effect using packages *survival*[48] and *coxme*[49]. To reflect latent class admixture proportions, we used proportional assignment: each individual was assigned to all of the 7 classes (6 LCA classes and the nulliparous) simultaneously with weights equal to class-specific posterior probabilities[50].

Study participants only entered the study provided they were alive in 1967, we therefore used 1967 as an entry date for left truncation in our model. Right-censored date of exit was either death, emigration or end of follow up, whichever came first. Two models were fitted: unadjusted model included only historical birth cohort and twin pair ID variables, whereas the adjusted model additionally included smoking, alcohol use, and BMI in 1975, and lifetime education (see variable description above). Statistical significant differences in mortality were defined by hazard ratios along with their 95% confidence intervals from the Cox proportional hazards model.

##### 3.3. Epigenetic aging

The differences in epigenetic age acceleration between the 6 latent reproductive classes and the nulliparous women were modeled in MPlus using the Bolck-Croon-Hagenaars (BCH) approach which controls for measurement error in the classification[55]. We used class-specific weights as training data to model the association between epigenetic age acceleration and latent classes. Similarly to survival analysis, we constructed two different models: an unadjusted model, which included independent variables for the twin pair ID and chronological age, and the adjusted model, which additionally included variables for smoking, alcohol use, BMI at the time of blood sampling, and lifetime education.

### Supplementary Results

Our sample consisted of 17,175 women with known live birth history. Extended Data Table 3 shows the sample distribution by zygosity, reproductive history, age and health-related variables for the data in the survival analysis (N=17,080), and for the subset with epigenetic age estimates (N=1,117). For all variables the number of observations is given (N) along with the number of individuals with missing data. There were 6,392 women that had not had childbirth during the study period, and the total average number of births per woman among the parous was 2.4 (SD 1.4). The average age at childbirth was 27.3 (SD 5.7), with 24.4 (SD 4.9) and 29.8 (SD 5.7) for first and last childbirth. The distribution of the ages at childbirth is illustrated in Extended Data Figure 6.

### Extended data

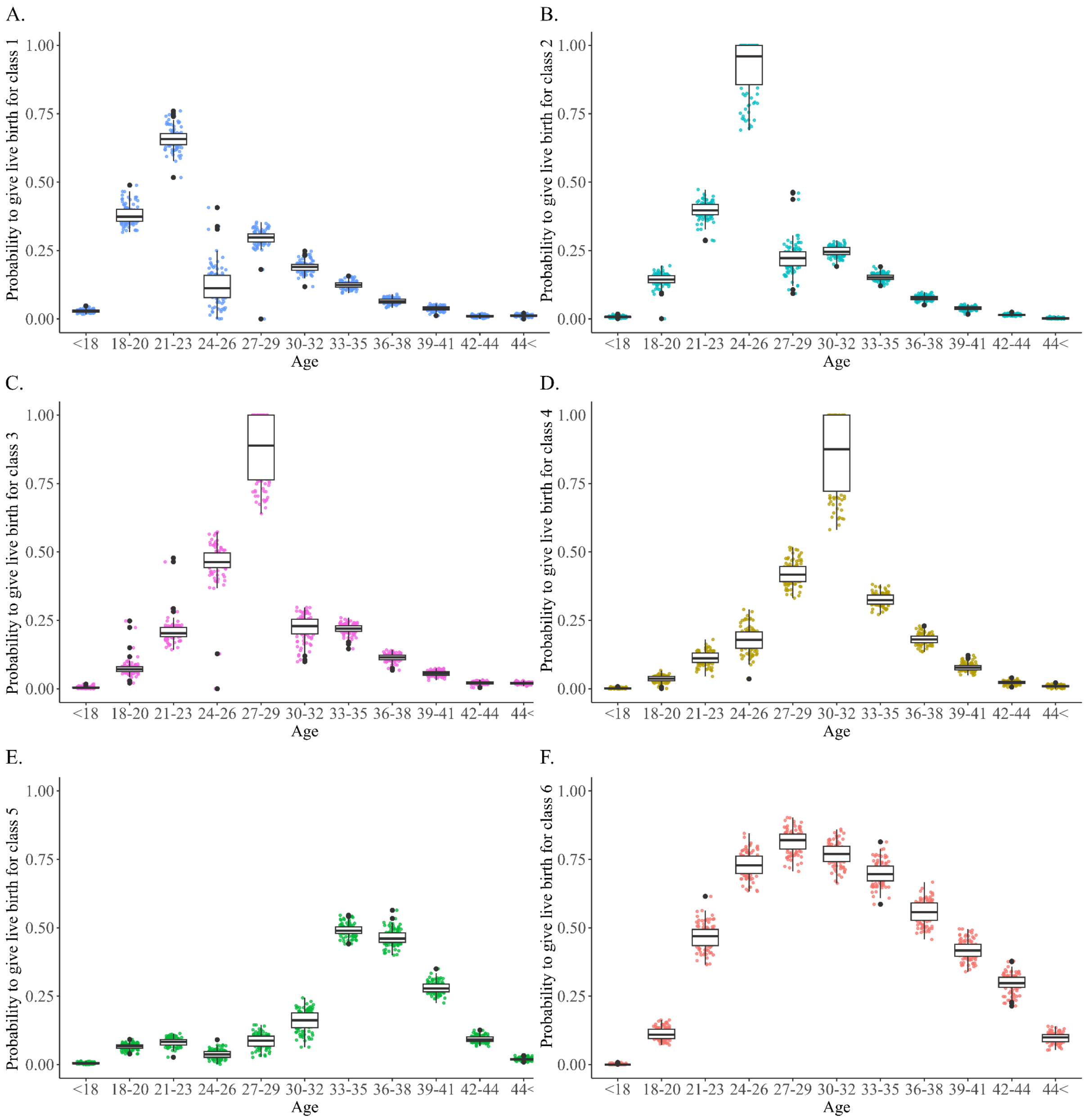


**Extended Data Figure 1. Probability to give birth at different ages from 100 parametric bootstrap rounds.** Jittered points represent probability estimates from the individual 100 bootstrap simulations. Figures 1A-F correspond to reproductive classes 1-6.

**Extended Data Table 1. Classification table of the latent class solution.** Classification probabilities for the most likely latent class (column) are given by latent class (row). The bolded diagonal represents the probability that an observation is part of a specific class that is classified as the same class.

| | Most likely latent class | | | | | |
|---|---|---|---|---|---|---|
| Latent class | 1 | 2 | 3 | 4 | 5 | 6 |
| 1 | **0.84** | 0.04 | 0.00 | 0.00 | 0.03 | 0.06 |
| 2 | 0.00 | **0.87** | 0.01 | 0.00 | 0.00 | 0.1 |
| 3 | 0.04 | 0.06 | **0.84** | 0.02 | 0.02 | 0.04 |
| 4 | 0.01 | 0.04 | 0.07 | **0.77** | 0.02 | 0.10 |
| 5 | 0.03 | 0.01 | 0.08 | 0.01 | **0.83** | 0.04 |
| 6 | 0.08 | 0.02 | 0.09 | 0.01 | 0.00 | **0.80** |

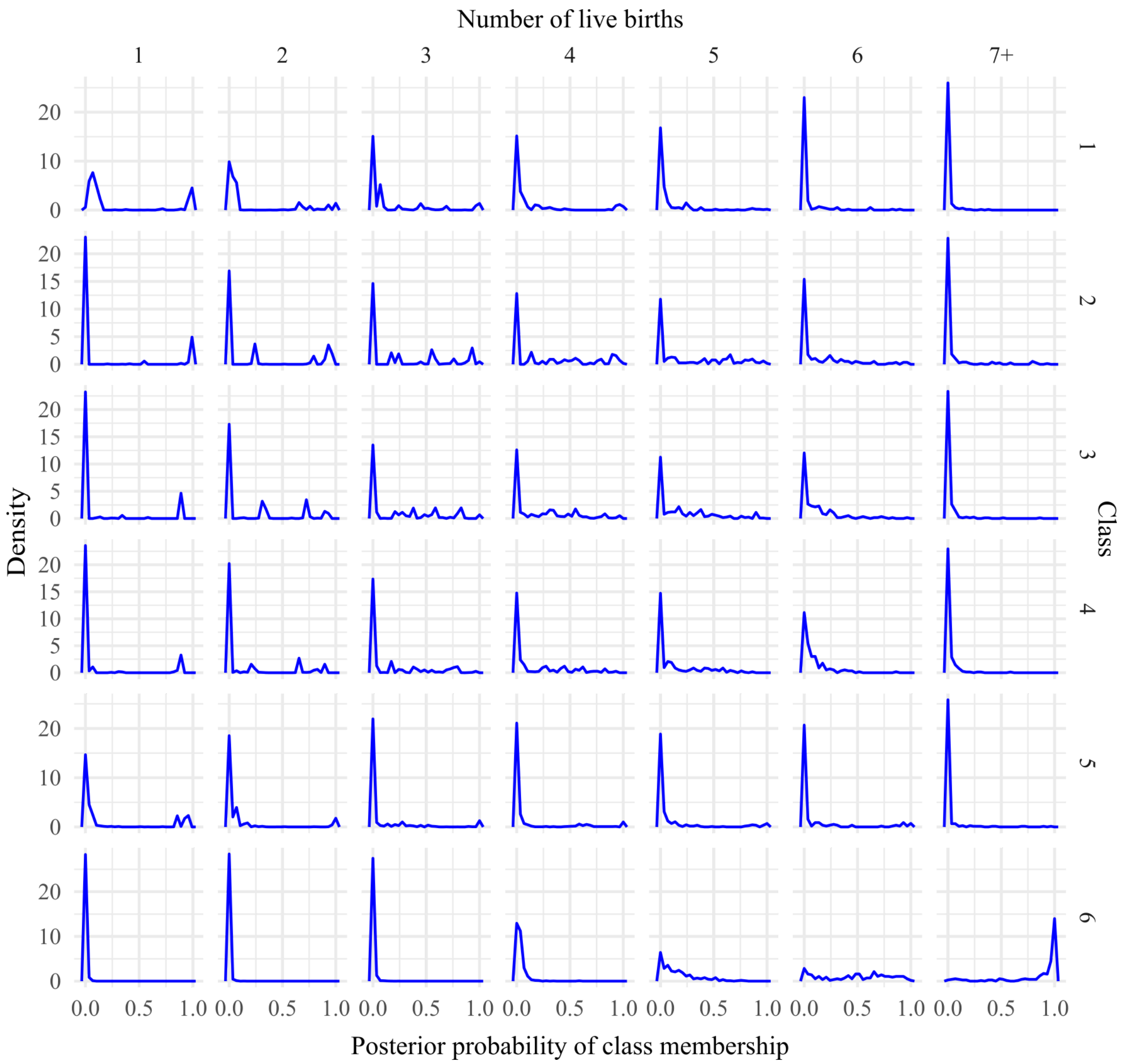


**Extended Data Figure 2. Number of pregnancies across reproductive classes.** Density plot of posterior probabilities for the membership of each class (rows) across women with different number of pregnancies across the women in the survival analysis from the LCA solution (N=10 688).

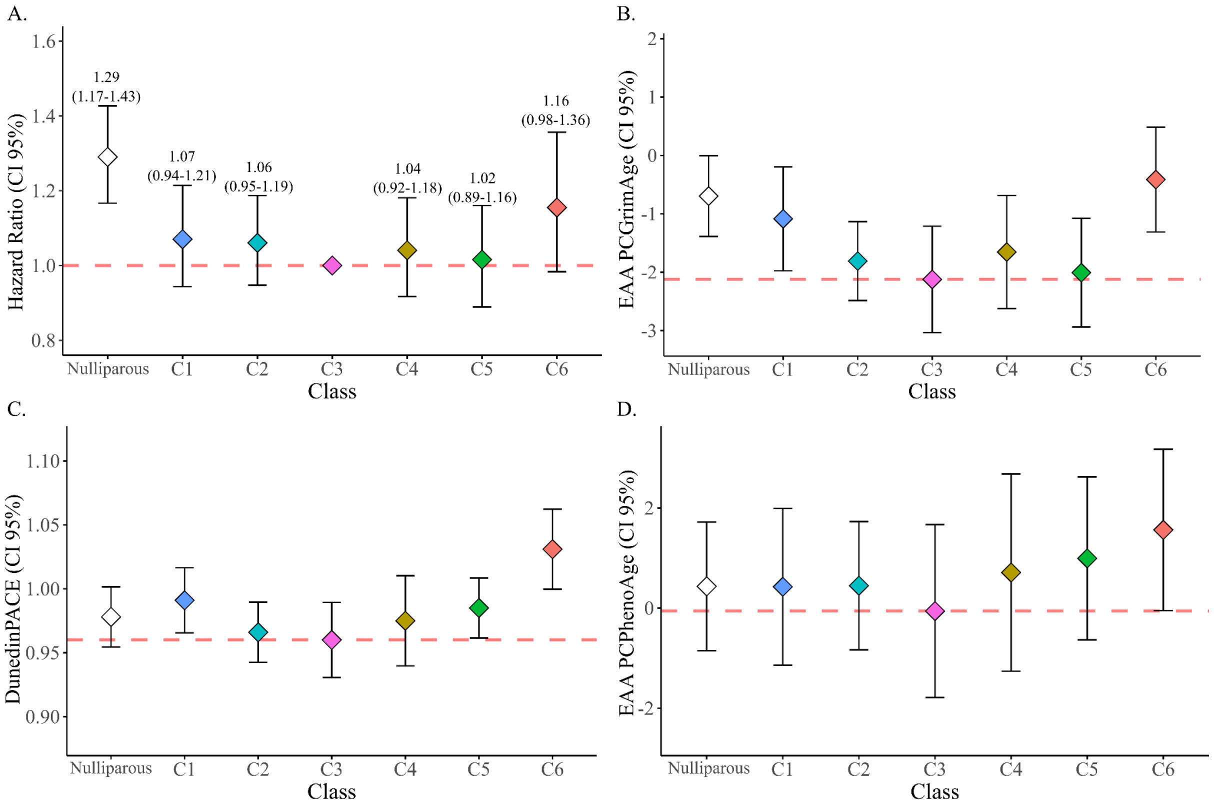


**Extended Data Figure 3. Models without covariate imputation.** A. Mortality hazard ratios with 95% confidence intervals for each latent class from a Cox proportional hazards model weighted with class-specific posterior propbabilities for the complete case sample (N=13,186). The red dashed line indicates the reference class in the survival model (class 3). The model is adjusted with relatedness, left truncation, alcohol use, smoking, bmi and education. B-D. Estimates for epigenetic age acceleration using PCGrimAge (B), PCPhenoAge (C) and DunedinPACE (D) with 95% confidence intervals for the model without covariate imputation using Full Information Maximum Likelihood (N=1117). The red dashed line is drawn for reference to contrast survival analysis. The epigenetic aging models are adjusted with chronological age, alcohol and tobacco use, and BMI at the time of blood sampling and lifetime education.

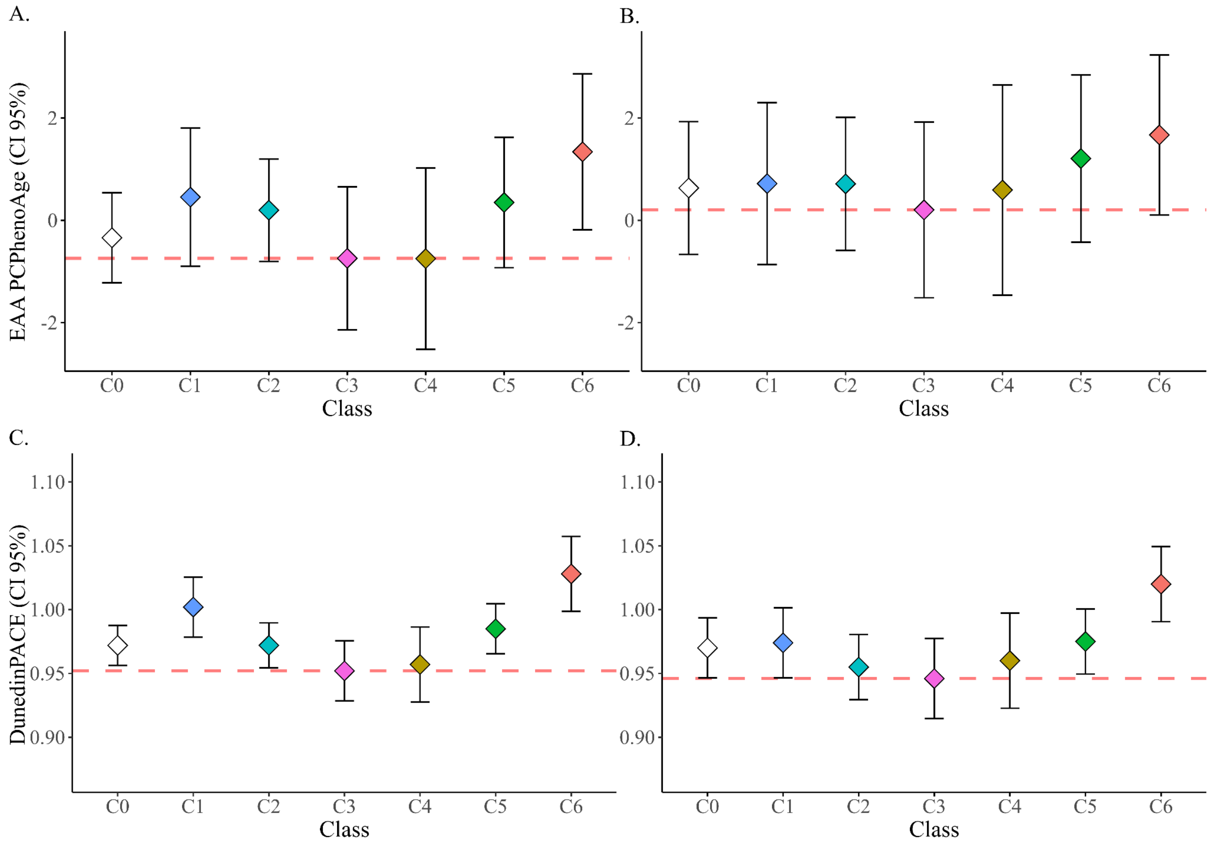


**Extended Data Figure 4. Epigenetic ages with PCPhenoAge and DunedinPACE.** A. Estimates and 95% confidence intervals for EAA by the PCPhenoAge clock for each class. The dashed red line is drawn for comparison to survival analysis at the estimates for class 3. B. Estimates and 95% confidence intervals for EAA by the PCPhenoAge clock for each class from a weighted Bolck-Croon-Hagenaars model, with additional adjustments for lifestyle factors. The dashed red line is drawn for comparison to survival analysis at the estimates for class 3. C. Estimates and 95% confidence intervals for EAA by the DunedinPACE clock for each class. The dashed red line is drawn for comparison to survival analysis at the estimates for class. B. Estimates and 95% confidence intervals for EAA by the DunedinPACE clock for each class from a weighted Bolck-Croon-Hagenaars model, with additional adjustments for lifestyle factors. The dashed red line is drawn for comparison to survival analysis at the estimates for class 3.

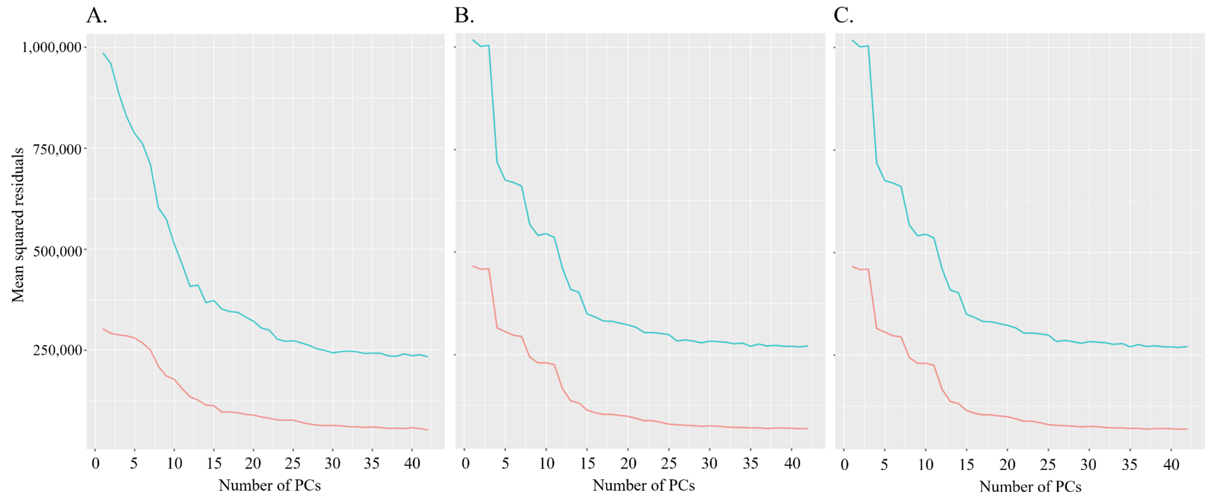


**Extended Data Figure 5. Mean squared residuals explained by the control probe principal components.**

The methylation data were preprocessed with 16, 15 and 16 for platforms 450k (A), EPICv1 (B), and EPICv2 (C), respectively.

**Extended Data Table 2. Comparison between all the fitted LCA models.** For each age year bin, classes 1–8 were fitted. Class proportions, entropy, average posterior probabilities for the most likely class (AvePP), sample size adjusted Bayesian information criterion (saBIC) and Akaike's information criterion (AIC) are given (for information criteria smaller is better). The final solution of 3-year age bins and 6 classes is bolded.

| Bin | Classes | Class proportions (%) | Entropy | AvePP | saBIC | AIC |
|---|---|---|---|---|---|---|
| 1 | 1 | | | | 169649 | 169374 |
| | 2 | 59.6, 40.4 | 0.44 | 0.84, 0.79 | 167895 | 167341 |
| | 3 | 61.7, 35.9, 2.4 | 0.66 | 0.86, 0.83, 0.77 | 167383 | 166549 |
| | 4 | 49.2, 38.8, 9.5, 2.5 | 0.64 | 0.86, 0.69, 0.68, 0.86 | 167213 | 166100 |
| | 5 | 50.7, 25.7,12.1, 8.7, 2.9 | 0.77 | 0.90, 0.80, 0.86, 0.73, 0.82 | 167220 | 165828 |
| | 6 | 44.2, 24.7, 10.3, 10.2, 7.7, 2.7 | 0.71 | 0.86, 0.70, 0.77, 0.66, 0.73, 0.84 | 167224 | 165553 |
| | 7 | 40.7, 22.8, 14.1, 10.2, 9.3, 2.0, 0.8 | 0.74 | 0.86, 0.69, 0.72, 0.93, 0.82, 0.83, 0.83 | 167338 | 165388 |
| | 8 | 37.8, 25.4, 14.5, 10.0, 9.3, 1.8, 0.6, 0.4 | 0.74 | 0.84, 0.71, 0.72, 0.75, 0.82, 0.83, 0.83, 0.81 | 167649 | 165419 |
| 2 | 1 | | | | 134826 | 134650 |
| | 2 | 58.9, 41.1 | 0.46 | 0.85, 0.81 | 132686 | 132329 |
| | 3 | 63.8, 33.7, 2.5 | 0.69 | 0.88, 0.78, 0.85 | 131903 | 131365 |
| | 4 | 47.3, 37.3, 12.8, 2.6 | 0.65 | 0.86, 0.71, 0.72, 0.86 | 131413 | 130694 |
| | 5 | 44.9, 19.5, 17.1, 15.4, 3.1 | 0.75 | 0.91, 0.83, 0.78, 0.78, 0.86 | 131244 | 130345 |
| | 6 | 42.8, 16.0, 15.2, 12.8, 10.0, 3.2 | 0.76 | 0.92, 0.77, 0.77, 0.84, 0.73, 0.92 | 131104 | 130024 |
| | 7 | 25.9, 17.8, 14.6, 14.6, 13.4, 10.1, 3.6 | 0.76 | 0.88, 0.80, 0.82, 0.76, 0.89, 0,76, 0.85 | 130939 | 129678 |
| | 8 | 28.0, 18.8, 16.0, 15.0, 9.5, 8.2, 3.9, 0.7 | 0.75 | 0.98, 0.75, 0.82, 0.77, 0.82, 0.82, 0.78, 0.87 | 131349 | 129908 |
| 3 | 1 | | | | 117058 | 116923 |
| | 2 | 61.5, 38.5 | 0.49 | 0.88, 0.78 | 114869 | 114594 |
| | 3 | 63.3, 34.5, 2.2 | 0.69 | 0.89, 0.78, 0.84 | 114257 | 113843 |
| | 4 | 46.9, 35.1, 15.3, 2.7 | 0.63 | 0.86, 0.74, 0.73, 0.84 | 113652 | 113098 |
| | 5 | 30.1, 22.8, 22.1, 21.9, 3.1 | 0.77 | 0.91, 0.75, 0.91, 0.89, 0.85 | 113186 | 112492 |
| | **6** | **26.1, 22.8, 20.3, 15.0, 12.5, 3.3** | **0.77** | **0.90, 0.77, 0.86, 0.76, 0.92, 0.83** | **112822** | **111989** |
| | 7 | 25.6, 21,5, 20.4, 15.1, 12.6, 3.8, 0.9 | 0.78 | 0.90, 0.76, 0.86, 0.77, 0.91, 0.72, 0.81 | 112899 | 111925 |
| | 8 | 25.3, 20.9, 20.5, 15.0, 12.3, 4.2, 0.9, 0.9 | 0.77 | 0.86, 0.77, 0.86, 0.76, 0.90, 0.65, 0.72, 0.80 | 113045 | 111932 |
| 4 | 1 | | | | 1011010 | 100995 |
| | 2 | 38.7, 61.3 | 0.46 | 0.77, 0,88 | 99225 | 98992 |
| | 3 | 38.2, 29.5, 32.3 | 0.57 | 0.73, 0.85, 0,1 | 98551 | 98197 |
| | 4 | 2.7, 40.4, 34.9, 21.9 | 0.69 | 0.81, 0.76, 0.89, 0.84 | 97983 | 97510 |
| | 5 | 20.1, 31.7, 21.1, 23.5, 3.7 | 0.81 | 0.97,0.85, 0.84, 0.89, 0.82 | 97253 | 96662 |
| | 6 | 3.6, 11.7, 9.3, 10.1, 28.4, 36.9 | 0.77 | 0.82, 0.84, 0.94, 0.87, 0.77, 0.78 | 96954 | 96243 |
| | 7 | 3.9, 10.3, 9.4, 35.3, 3.2, 16.5, 21.3 | 0.77 | 0.82, 0.82, 0.76, 0.81, 0.90, 0.78, 0.77 | 96789 | 95959 |
| | 8 | 10.1, 4.2, 1.2, 15.4, 3.7, 30.0, 15.7, 19.9 | 0.76 | 0.78, 0.74, 0.85, 0.84, 0.87, 0.78, 0.71, 0.80 | 96797 | 95848 |

**Extended Data Table 3. Descriptive variables of the participants in the survival and epigenetic age acceleration analyses.** Levels, level-specific sample numbers with corresponding proportions are given for categorical variables whereas range, mean and standard deviation (SD) are given for continuous variables. For all variables the number of observations is given (N) along with the number of individuals with missing data (NA(%)). Zygosities for all twins are coded as monozygotic (MZ), dizygotic (DZ) and unknown (XZ), who are still confirmed as twins. For survival data, BMI, smoking and alcohol use are reported for the start of the follow-up in 1975, whereas for epigenetic aging these covariates are reported for the time of blood sampling.

Education is reported as lifetime education from the questionnaire in 1975 or 1981 whichever was the most recent for any participant. *Means and standard deviations for ages at childbirth and number of births are calculated for parous women only.

| | Survival | | | | Epigenetic age acceleration | | | |
|---|---|---|---|---|---|---|---|---|
| | N | Category/ Range | Prop(%)/ Mean(SD) | NA(%) | N | Category/ Range | Prop(%)/ Mean(SD) | NA(%) |
| **Zygosity** | 17 080 | | | | 1 117 | | | |
| | 4 221 | MZ | 24.7 | | 500 | MZ | 44.8 | |
| | 11 663 | DZ | 75.7 | | 614 | DZ | 55.0 | |
| | 1 134 | XZ | 6.6 | | 3 | XZ | 0.2 | |
| **Pregnancies*** | 10 688 | 1-15 | 2.4 (1.4) | | 855 | 1-12 | 2.6 (1.8) | |
| **Pregnancies** | | | | | | | | |
| | 6 392 | 0 | 37.4 | | 262 | 0 | 23.5 | |
| | 2 603 | 1 | 15.2 | | 219 | 1 | 19.6 | |
| | 4 250 | 2 | 24.9 | | 310 | 2 | 27.8 | |
| | 2 273 | 3 | 13.3 | | 157 | 3 | 14.1 | |
| | 1 562 | 3< | 9.1 | | 169 | 3< | 15.1 | |
| **Age at 1st birth*** | 10 688 | 13-44 | 25.4 (4.9) | | 855 | 17-44 | 25.9 (5.1) | |
| **Age at last birth*** | 10 688 | 16-50 | 30.8 (5.7) | | 855 | 17-47 | 31.1 (5.7) | |
| **Age at death** | 5 588 | 40-107 | 76.6 (13.1) | | 300 | 46-99 | 82.8 (8.9) | |
| **Age in 1975** | 17 080 | 17-95 | 33.1 (14.9) | | 1 117 | 17-65 | 36.1 (10.8) | |
| **Historical Cohort** | 17 080 | | | | 1 117 | | | |
| | 101 | Industrial | 0.6 | | 0 | Industrial | 0 | |
| | 1 561 | Pre-indep. | 9.1 | | 18 | Pre-indep. | 1.6 | |
| | 3 824 | Post-indep. | 22.4 | | 604 | Post-indep. | 54.1 | |
| | 2 031 | WW II | 11.9 | | 125 | WW II | 11.2 | |
| | 9 563 | Cold War | 56.0 | | 370 | Cold War | 33.1 | |
| **BMI** | 13 272 | 14.5-50.2 | 22.7 (3.6) | 3 808 (22.3) | 1008 | 16.9-47.7 | 26.9 (4.9) | 109 (9.8) |
| **Smoking** | 13 352 | | | 3 728 (21.8) | 966 | | | 151 (13.5) |
| | 8 531 | Never | 63.9 | | 667 | Never | 69.0 | |
| | 348 | Former | 2.6 | | 169 | Former | 17.1 | |
| | 1 371 | Infrequent | 10.3 | | 130 | Current | 13.5 | |
| | 1 430 | Light | 10.7 | | | | | |
| | 1 318 | Medium | 9.9 | | | | | |
| | 354 | Heavy | 2.7 | | | | | |
| **Alcohol** | 13 359 | | | 3 721 (21.8) | 898 | | | 219 (19.6) |
| | 3 529 | Abstainer | 26.4 | | 301 | Non-drinker | 33.5 | |
| | 91 | Former | 0.7 | | 59 | Infrequent | 6.6 | |
| | 1 308 | Infrequent | 9.8 | | 538 | Frequent | 59.9 | |
| | 8 205 | Low | 61.4 | | | | | |
| | 226 | High | 1.7 | | | | | |
| **Education** | 13 699 | | | 3 381 (19.8) | 1 117 | | | 0 |
| | 6 277 | Primary | 45.8 | | 428 | Primary | 38.3 | |
| | 5 330 | Secondary | 38.9 | | 455 | Secondary | 40.7 | |
| | 2 092 | Tertiary | 15.3 | | 234 | Tertiary | 20.9 | |

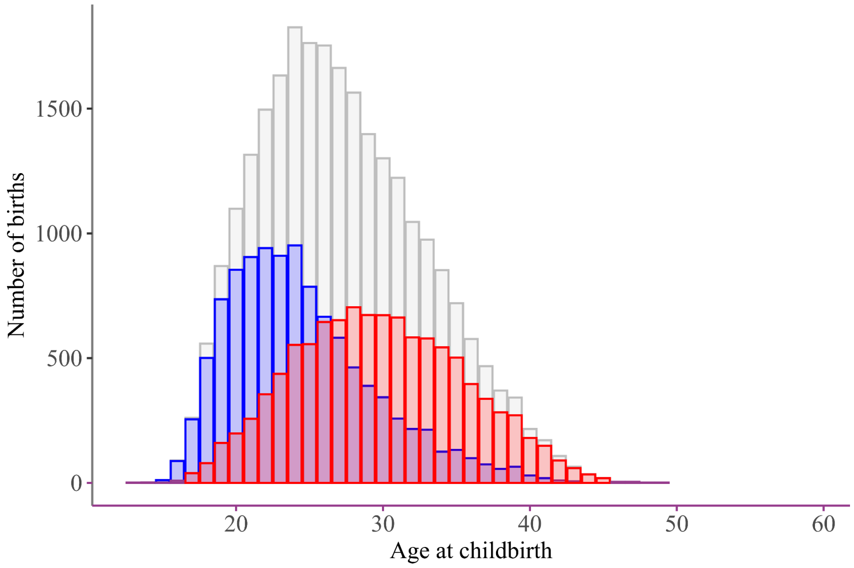


**Extended Data Figure 6. Number of live births by maternal age.** All births (N=25 801) are given as gray bars, blue bars represent only first births (N=10 688) and red bars indicate last births (N=10 688).

**Extended Data Table 4. Cross-class comparisons from BCH models of epigenetic age acceleration.** The comparisons are given for PCGrimAge, PCPhenoAge and DunedinPACE. Estimates for differences between classes are given with their respective standard errors (SE) and p-values are given for models 1(unadjusted) and 2 (adjusted for BMI, smoking, alcohol and education). Significant ($p<0.05$) differences are bolded.

| **PCGrimAge** | | Model 1 | | | Model 2 | |
|---|---|---|---|---|---|---|
| Comparison | Estimate | SE | p-value | Estimate | SE | p-value |
| C1-C0 | 0.331 | 0.514 | 0.52 | -0.268 | 0.475 | 0.573 |
| C1-C3 | **1.534** | **0.61** | **0.012** | 1.085 | 0.573 | 0.058 |
| C2-C0 | **-0.889** | **0.385** | **0.021** | **-1.053** | **0.366** | **0.004** |
| C2-C1 | **-1.22** | **0.515** | **0.018** | -0.786 | 0.441 | 0.075 |
| C2-C3 | 0.314 | 0.513 | 0.541 | 0.3 | 0.476 | 0.529 |
| C3-C0 | **-1.203** | **0.473** | **0.011** | **-1.353** | **0.453** | **0.003** |
| C4-C0 | **-1.443** | **0.56** | **0.01** | **-1.269** | **0.524** | **0.016** |
| C4-C1 | **-1.774** | **0.65** | **0.006** | -1.001 | 0.622 | 0.108 |
| C4-C2 | -0.554 | 0.557 | 0.32 | -0.216 | 0.528 | 0.683 |
| C4-C3 | -0.24 | 0.651 | 0.712 | 0.084 | 0.628 | 0.893 |
| C4-C5 | -0.541 | 0.67 | 0.42 | 0.005 | 0.624 | 0.994 |
| C5-C0 | -0.903 | 0.504 | 0.073 | **-1.273** | **0.475** | **0.007** |
| C5-C1 | **-1.233** | **0.608** | **0.042** | -1.006 | 0.565 | 0.075 |
| C5-C2 | -0.013 | 0.503 | 0.979 | -0.22 | 0.458 | 0.631 |
| C5-C3 | 0.301 | 0.565 | 0.595 | 0.08 | 0.537 | 0.882 |
| C6-C0 | -0.403 | 0.498 | 0.419 | 0.086 | 0.528 | 0.87 |
| C6-C1 | -0.733 | 0.601 | 0.223 | 0.354 | 0.596 | 0.552 |
| C6-C2 | 0.487 | 0.513 | 0.342 | **1.14** | **0.54** | **0.035** |
| C6-C3 | 0.801 | 0.6 | 0.182 | **1.439** | **0.634** | **0.023** |
| C6-C4 | 1.041 | 0.668 | 0.119 | 1.355 | 0.719 | 0.059 |
| C6-C5 | 0.5 | 0.612 | 0.414 | **1.36** | **0.638** | **0.033** |

| **PCPhenoAge** | Model 1 | | | Model 2 | | |
|---|---|---|---|---|---|---|
| Comparison | Estimate | SE | p-value | Estimate | SE | p-value |
| C1-C0 | 0.795 | 0.813 | 0.328 | 0.086 | 0.861 | 0.92 |
| C1-C3 | 1.198 | 1.054 | 0.256 | 0.515 | 1.076 | 0.633 |
| C2-C0 | 0.539 | 0.673 | 0.423 | 0.082 | 0.697 | 0.907 |
| C2-C1 | -0.256 | 0.854 | 0.765 | -0.004 | 0.849 | 0.996 |
| C2-C3 | 0.942 | 0.911 | 0.301 | 0.51 | 0.919 | 0.578 |
| C3-C0 | -0.403 | 0.812 | 0.62 | -0.429 | 0.813 | 0.598 |
| C4-C0 | -0.408 | 0.992 | 0.681 | -0.039 | 0.968 | 0.968 |
| C4-C1 | -1.203 | 1.158 | 0.299 | -0.125 | 1.214 | 0.918 |
| C4-C2 | -0.947 | 1.06 | 0.372 | -0.121 | 1.075 | 0.911 |
| C4-C3 | -0.005 | 1.227 | 0.997 | 0.39 | 1.219 | 0.749 |
| C4-C5 | -1.097 | 1.142 | 0.337 | -0.616 | 1.14 | 0.589 |
| C5-C0 | 0.689 | 0.792 | 0.384 | 0.576 | 0.799 | 0.47 |
| C5-C1 | -0.105 | 0.957 | 0.912 | 0.49 | 0.982 | 0.618 |
| C5-C2 | 0.15 | 0.823 | 0.855 | 0.495 | 0.837 | 0.554 |
| C5-C3 | 1.092 | 0.965 | 0.258 | 1.005 | 0.972 | 0.301 |
| C6-C0 | 1.681 | 0.893 | 0.06 | 1.038 | 0.977 | 0.288 |
| C6-C1 | 0.886 | 1.042 | 0.396 | 0.952 | 1.107 | 0.39 |
| C6-C2 | 1.141 | 0.932 | 0.221 | 0.956 | 1 | 0.339 |
| C6-C3 | 2.083 | 1.101 | 0.059 | 1.467 | 1.181 | 0.214 |
| C6-C4 | 2.089 | 1.263 | 0.098 | 1.077 | 1.342 | 0.422 |
| C6-C5 | 0.991 | 1.024 | 0.333 | 0.461 | 1.11 | 0.678 |
| **PACE** | Model 1 | | | Model 2 | | |
| Comparison | Estimate | SE | p-value | Estimate | SE | p-value |
| C1-C0 | **0.03** | **0.015** | **0.042** | 0.004 | 0.015 | 0.779 |
| C1-C3 | **0.05** | **0.018** | **0.006** | 0.028 | 0.018 | 0.131 |
| C2-C0 | 0 | 0.012 | 0.993 | -0.015 | 0.012 | 0.216 |
| C2-C1 | **-0.03** | **0.015** | **0.048** | -0.019 | 0.014 | 0.19 |
| C2-C3 | 0.02 | 0.016 | 0.223 | 0.009 | 0.016 | 0.57 |
| C3-C0 | -0.02 | 0.015 | 0.175 | -0.023 | 0.014 | 0.102 |
| C4-C0 | -0.015 | 0.017 | 0.381 | -0.009 | 0.017 | 0.58 |
| C4-C1 | **-0.046** | **0.02** | **0.021** | -0.013 | 0.02 | 0.508 |
| C4-C2 | -0.015 | 0.018 | 0.4 | 0.005 | 0.018 | 0.767 |
| C4-C3 | 0.005 | 0.021 | 0.83 | 0.014 | 0.021 | 0.499 |
| C4-C5 | -0.028 | 0.018 | 0.132 | -0.014 | 0.018 | 0.426 |
| C5-C0 | 0.012 | 0.013 | 0.321 | 0.005 | 0.012 | 0.675 |
| C5-C1 | -0.018 | 0.016 | 0.252 | 0.001 | 0.015 | 0.959 |
| C5-C2 | 0.013 | 0.013 | 0.347 | 0.019 | 0.012 | 0.114 |
| C5-C3 | **0.032** | **0.015** | **0.034** | 0.028 | 0.015 | 0.056 |
| C6-C0 | **0.056** | **0.017** | **0.001** | **0.05** | **0.018** | **0.005** |
| C6-C1 | 0.025 | 0.019 | 0.184 | **0.046** | **0.02** | **0.019** |
| C6-C2 | **0.056** | **0.017** | **0.001** | **0.065** | **0.019** | **0** |
| C6-C3 | **0.075** | **0.02** | **0** | **0.074** | **0.022** | **0.001** |
| C6-C4 | **0.071** | **0.022** | **0.001** | **0.06** | **0.024** | **0.012** |
| C6-C5 | **0.043** | **0.018** | **0.015** | **0.045** | **0.019** | **0.016** |
